# A General Transformer-Based Multi-Task Learning Framework for Predicting Interaction Types between Enzyme and Small Molecule

**DOI:** 10.1101/2025.10.09.681419

**Authors:** Vahid Atabaigi Elmi, Roman Joeres, Olga V. Kalinina

## Abstract

Predicting enzyme–small molecule interactions is critical for drug discovery and generally understanding the biochemical processes of life. While recent deep learning approaches have shown promising results, several challenges remain: the lack of a comprehensive training dataset, architectures lacking communication between representations of enzymes and small molecules, the tendency to simplify the problem as enzyme–substrate vs. enzyme—non-interacting, and thereby, misclassify enzyme–inhibitor pairs as substrates, and the negligence of the true impact of data leakage on the model’s performance. To address these issues, we present EMMA (**E**nzyme–small **M**olecule interaction **M**ulti-head **A**ttention), a transformer-based multi-task learning framework designed to learn pairwise interaction signals between enzymes and small molecules, that can well generalize to out-of-distribution data. EMMA operates directly on the SMILES string representation of small molecules and enzyme sequences, with two classification heads that distinguish enzyme—non-interacting, enzyme—substrate, and enzyme—inhibitor pairs. By evaluating EMMA under five distinct data-splitting regimes that control for different types of data leakage, we demonstrate that EMMA achieves a strong and robust performance, particularly for previously unseen combinations of enzymes and small molecules. Further, a deeper analysis highlights that the topological properties of the enzyme—small molecule interaction network are crucial for the model performance and its ability to generalize, yet again stressing the decisive role of well-designed training datasets for successful model training.

## 1 Main

Enzymes are proteins that catalyze essential biological reactions, playing key roles in metabolism, energy transfer, and signal transduction [1]. Understanding enzyme— small molecule interactions is fundamental in biology and drug design. These interactions can be categorized into three types: (i) enzyme—substrate, when a small molecule binds to the enzyme’s active site and is converted into a product, (ii) enzyme—non-interacting, where a small molecule with very low affinity or incompatibility for the enzyme cannot bind to the active site, and (iii) enzyme—regulator pairs, where small molecules such as inhibitors or activators modulate enzymatic activity [1].

The functions of many enzymes are still poorly understood, as only about 3% of human proteins have high-quality functional annotations supported by experimental evidence [2]. Computational methods, particularly machine learning, have revolutionized biological research by enabling large-scale data analysis and uncovering insights that were previously unattainable through manual methods [3]. However, predicting enzyme—small molecule interactions remains a challenge due to their larger complexity compared even to drug—target interactions. This complexity stems from the high specificity of enzymes, dynamic and transient interactions with ligands, and conformational flexibility during catalysis [4–6]. Additionally, enzymes are often multifunctional and promiscuous binders [7, 8]. The evolutionary diversity of enzymes and the lack of high-quality data exacerbate these challenges [5, 6], making the design of efficient computational methods to predict enzyme functionality even more challenging.

Computational methods for predicting interactions of enzymes and small molecules can be docking-based [9–12] and non-docking [13–17] (ligand-based or data-driven), each with its own trade-offs in accuracy, interpretability, and scalability. Since docking-based methods are prohibitively slow for large datasets [18], this work addresses non-docking prediction, which is typically done by means of machine learning, or more precisely, nowadays deep learning. In such non-docking methods, training a model to predict all interaction types (e.g., substrates, regulators, and non-interactors) remains challenging due to the lack of comprehensive and curated enzyme—small molecule datasets. Consequently, many studies simplify the task to a binary classification problem, distinguishing known enzyme—substrate pairs from either experimentally validated [13] or synthetically generated [14–17, 19] non-substrates. However, the term *non-substrate* can refer to both small molecules that fail to bind an enzyme due to low affinity or structural incompatibility, as well as to tightly binding regulating ligands (inhibitors or activators). We use the term *non-interacting small molecule* to avoid this confusion.

Since experimental data on non-interacting ligands is scarce, synthetically generated enzyme—non-interacting pairs are often used to train computational models, which may introduce biases and data shifts caused by the generation process. Furthermore, simplification to a binary classification task introduces a critical limitation: the model is trained to differentiate high-affinity enzyme—substrate interactions from presumed enzyme—non-interacting pairs, and thus is prone to misclassifying other high-affinity binders such as inhibitors or cofactors as substrates, leading to incorrect predictions. Furthermore, many datasets are skewed towards interactions with energy-transfer small molecules (e.g., the dataset from [14] contains 1,379 small molecules in total and ~ 28% of interactions involve an energy-transfer small molecule). This imbalance makes these interactions easily predictable, often inflating model performance metrics, but they are less informative for truly learning interaction patterns with diverse substrate molecules.

Another cause of artificially overinflated performance in models for enzyme— small molecule interaction predictions that has to be accounted for is data leakage, in particular inter-sample similarity leakage that occurs when similar data points appear in both training and test sets [20]. To minimize the impact of data leakage in biological datasets and ensure robust model evaluation, various tools have been developed, including DeepChem [3], LoHi-Splitter [21], GraphPart [22], and DataSAIL [23]. Among these tools, DataSAIL is the only one capable of minimizing inter-sample similarity leakage in two-dimensional datasets, such as the enzyme—small molecule interactions.

In this study, we present a comprehensive framework for enzyme—small molecule interaction prediction that addresses these challenges through three key contributions: (i) we constructed the novel EMMI (Enzyme—sMall Molecule Interaction) dataset encompassing experimentally confirmed enzyme—substrate, enzyme—non-interacting, and enzyme—inhibitor pairs, significantly expanding beyond previous binary formulations; (ii) we implemented rigorous data splitting protocols using DataSAIL to control inter-sample similarity leakage between training and test sets; and (iii) we developed a transformer-based multi-task learning architecture that processes both enzyme and small molecule features. The model employs two classification heads: an interaction head, which distinguishes between low- and high-affinity interactions, and a subclass head, which specifies the interaction type: either inhibitor or substrate. Given the lack of comparable models, we benchmarked EMMA against random forests and demonstrated its effectiveness in distinguishing between enzyme—substrate, enzyme— non-interacting, and enzyme—inhibitor pairs, while maintaining robustness against data leakage.

## 2 Results

### 2.1 The EMMI dataset

We created the EMMI (Enzyme-sMall Molecule Interactions) dataset comprising a total of 147,224 samples, divided into three types: enzyme—non-interacting, enzyme— substrate, and enzyme—inhibitor pairs. The dataset comprises 15,652 unique enzymes and 28,977 unique small molecules, resulting in an overall enzyme-to-small molecule ratio of 0.54 (Table 1). The dataset is well-balanced in size across all enzyme—small molecule relation types, while still capturing diversity in both enzyme and small molecule entries. We intentionally kept the number of enzyme—non-interacting pairs (79,668 pairs) higher than the combined number of enzyme—substrate and enzyme— inhibitor pairs (67,556 pairs) to better reflect the biological setting, where most enzymes do not interact with the majority of small molecules. The ratio of enzyme— substrate to enzyme—inhibitor samples is ~ 1. The higher enzyme-to-small molecule ratio in the enzyme—substrate subset highlights its enzyme-centric nature, whereas the enzyme—non-interacting and —inhibitor subsets are more small molecule-rich.

**Table 1.** EMMI dataset statistics and composition after and (before) clustering-based down-sampling. Enz/mol: enzyme /molecule; enz: enzyme; NI: non-interacting; sub: substrate; inh: inhibitor

| Dataset | samples | Unique entries |  | Enz/mol ratio |
| --- | --- | --- | --- | --- |
|  |  | Enzyme ID | molecule ID |  |
| Enz-NI | 79,668 (319,413) | 4,351 (4,351) | 14,283 (164,317) | 0.305 (0.027) |
| Enz-sub | 34,221 (34,221) | 11,343 (11,343) | 6,569 (6,569) | 1.730 (1.730) |
| Enz-inh | 33,335 (807,173) | 5,997 (5,997) | 10,436 (564,613) | 0.575 (0.011) |
| Subclass | 67,556 (841,394) | 17,340 (17,340) | 17,005 (571,182) | 1.020 (0.030) |
| Interaction | 147,224 (1,161,365) | 15,652 (15,652) | 28,977 (691,959) | 0.540 (0.023) |

A single enzyme and a single small molecule can be a member in the pairs with different labels (substrate, inhibitor, or non-interacting), but the distribution of these labels among enzymes and small molecules is markedly different in the EMMI dataset (Figure 1). The enzyme distribution reveals substantial overlap between substrate-binding and inhibitor-binding enzymes: while 8,104 enzymes are uniquely substrate-binding, 1,630 are uniquely inhibitor-binding, and 1,145 are uniquely non-interacting, there are 4,773 multi-label enzymes with 1,266 of them having three labels, reflecting the multifunctional nature of enzymes [24, 25]. In contrast, the distribution of small molecules shows far less overlap: with 1,966 multi-label small molecules and only 349 small molecules being simultaneously labeled as substrates, inhibitors, and non-interacting. Most small molecules are strongly partitioned into either substrates (5,586), inhibitors (8,852), or non-interacting (12,518), consistent with their specialized roles towards enzymes. Collectively, these results highlight an important asymmetry; enzymes are often multifunctional and promiscuous binders [7], while small molecules tend to exhibit more exclusive binding roles due to their intrinsic chemical specificity and the selective optimization processes of drug discovery, which deliberately favor narrow, well-defined interactions to maximize efficacy and minimize off-target effects [26–28]. (Figure 1).

**Fig. 1.**
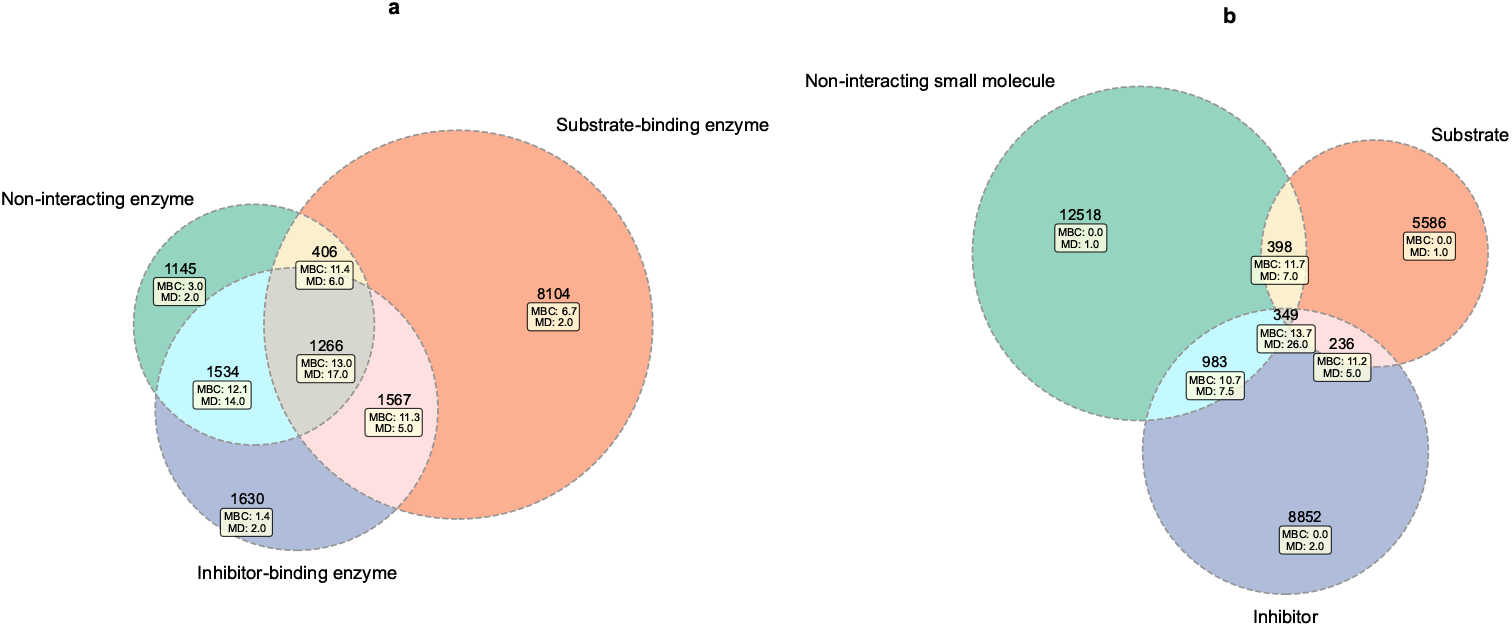
Distribution of class labels in the EMMI dataset for enzymes **(a)** and small molecules **(b)**. MBC: median betweenness centrality. MD: median degree. Note: Both MBC (log+1) and MD are log-scaled.

### 2.2 Types of inter-sample similarity leakage

We propose that inter-sample similarity leakage can be further partitioned into two subtypes, each affecting the model’s performance differently. Before defining these subtypes, it is helpful to represent the enzyme–small molecule interaction network as a bipartite graph, where enzymes and small molecules form two disjoint node (vertex) sets, and edges between node sets represent observed interactions. When this network is split into training and test sets based on either node type (enzyme or small molecule), inter-sample similarity within the vertex sets can cause information leakage in two forms.

The first form is *single-label inter-sample similarity leakage* (SLSL). This happens when similar data points—e.g., similar enzymes (or their close variants) interact with multiple small molecules—appear in both the training and test sets with the same label (Figure 2 a). This form of leakage can artificially boost a model’s performance: after a few training iterations, the model may effectively memorize the features of the bridging enzymes, leveraging their repeated presence across multiple substrates rather than learning genuine interaction patterns—in other words, relying on correlation through repetition rather than true causality. Theoretically, single-label nodes with high connectivity—in terms of degree or betweenness centrality (BC)—that bridge the training and test sets are more likely to mislead the model toward memorization (Figure 2 a vs. c), due to the dominant availability of their features during training (Figure 2 e vs. g). The impact of SLSL can be more severe in tree-based models, such as random forests and XGBoost, where feature importance is often ranked based on how frequently features are used to create decision splits.

**Fig. 2.**
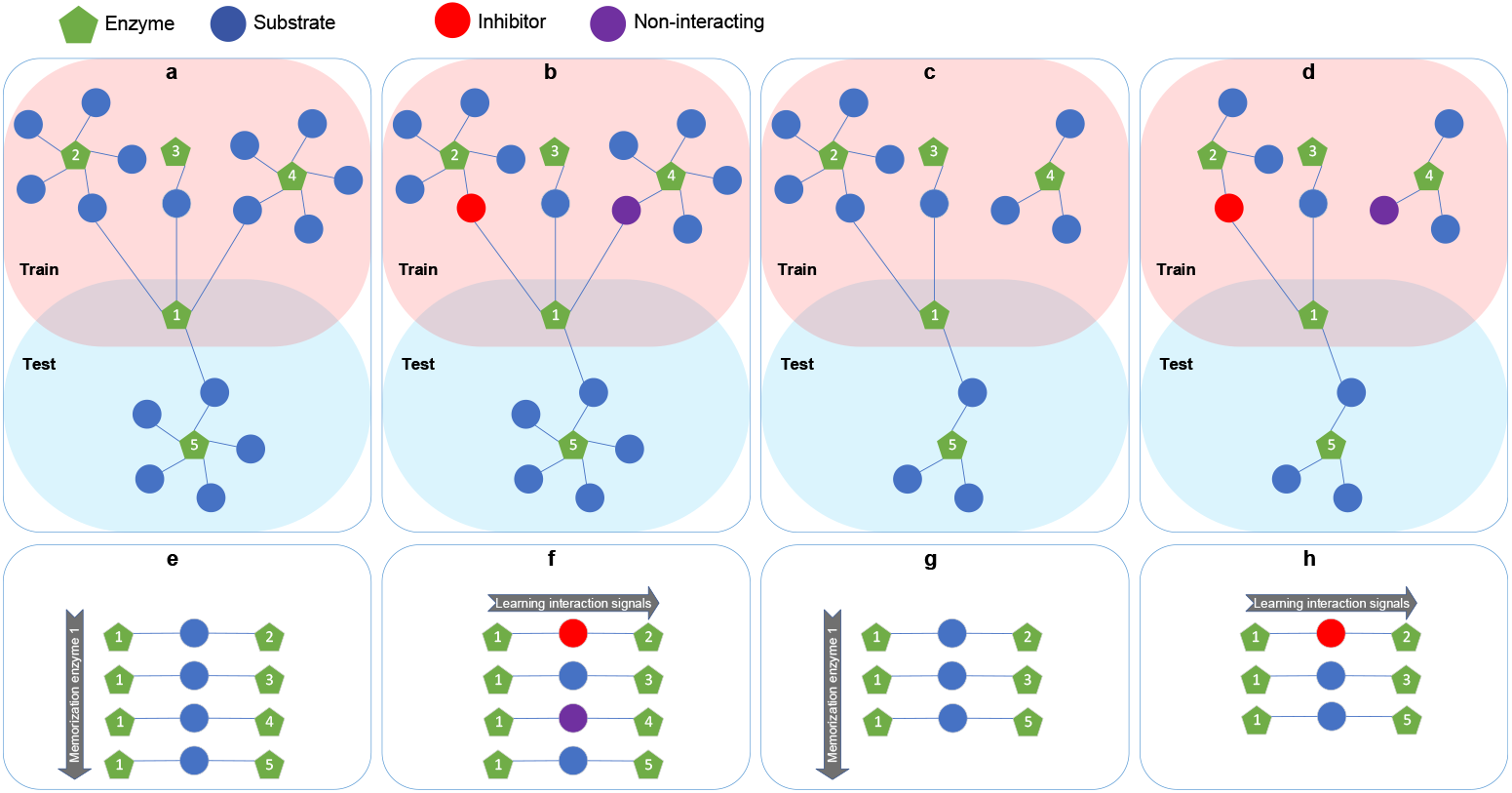
Different forms of inter-sample similarity leakage across training and test sets. **(a, c)**Single-label inter-sample similarity leakage (SLSL) where single-label enzyme with high and low connectivity (degree and betweenness centrality) shared between training and test sets. **(b, d)** Multi-label inter-sample similarity leakage (MLSL) where multi-label enzyme with high and low connectivity shared between training and test sets. **(e, f, g, h)** Frequent occurrence of enzyme 1 features under high and low connectivity and its impact on model learning ability.

The second form is *multi-label inter-sample similarity leakage* (MLSL). This occurs when the same node or very similar nodes are present in both the training and test sets but are associated with different labels (Figure 2 b and d). This form of leakage can negatively affect a model’s performance, particularly if the model tends to memorize rather than learn interaction signals. However, these multi-label nodes are more likely to lead the model toward learning interaction signals if they have low to moderate connectivity. In contrast, when these nodes are highly connected, the model is more likely to memorize the shared node across many interactions, because it is present in many sample pairs during the training process. If this node (or its close variants) is labeled differently in the test set, the model is likely to fail. Unlike single-label nodes, multi-label nodes tend to have a higher median BC and degree (Figure 1).

Overall, we believe that the tendency of the model to memorize or learn the interaction signals is not only related to the magnitude and the form of the inter-sample similarity leakage, but also the topology of the enzyme—small molecule interaction network in terms of connectivity of nodes in the training and test sets (Figure 2 b vs. d and f vs. h); single-label nodes with high connectivity are more prone to being memorized by the model due to the dominant availability of their features during training, whereas multi-label nodes with moderate connectivity can trigger the model to learn interaction signals.

### 2.3 Data split regimes

To assess model performance, we employed four one-dimensional split regimes (random, enzyme-based, small molecule-based, and label-based) as well as a two-dimensional split regime (Section 4.1.5, Table 2), ensuring an 80:20 % ratio between training and test sets in each case. For all regimes, we kept the ratio of enzyme— substrate to enzyme—inhibitor pairs close to 1 and the ratio of enzyme—interacting (enzyme—substrate or enzyme—inhibitor) to enzyme—non-interacting pairs ~ 0.43. Each one-dimensional split produces training and test sets of ~ 118K and ~ 29K, respectively, retaining all the samples from the EMMI dataset, and the two-dimensional split yields a smaller total set due to its stricter partitioning criteria: ~65K in the training and ~16K in the test sets.

**Table 2.** Overview of different splits with corresponding sample and label ratios. Enz–Inh: Enzyme–inhibitor. Enz–Sub: Enzyme–substrate. Enz–NI: Enzyme–non-interacting.

| Split | Num. Samples |  | Enz-Inh/Enz-Sub |  | Enz-Sub/Enz-NI |  | Enz-Inh/Enz-NI |  |
| --- | --- | --- | --- | --- | --- | --- | --- | --- |
|  | Training | Test | Training | Test | Training | Test | Training | Test |
| Random | 117,779 | 29,445 | 0.974 | 0.974 | 0.430 | 0.430 | 0.418 | 0.418 |
| Enzyme-based | 117,771 | 29,453 | 0.985 | 0.932 | 0.424 | 0.454 | 0.417 | 0.423 |
| Small molecule-based | 117,863 | 29,361 | 1.033 | 0.786 | 0.403 | 0.547 | 0.416 | 0.430 |
| Label-based | 117,594 | 29,630 | 0.973 | 0.977 | 0.433 | 0.417 | 0.421 | 0.408 |
| Two-dimensional | 64,881 | 16,083 | 1.193 | 0.763 | 0.324 | 0.666 | 0.387 | 0.508 |

Each split method exhibits distinct characteristics with respect to inter-sample similarity leakage (see Table 3 and Section 4.1.5 for details). In the random split, both single- and multi-label nodes are distributed arbitrarily across train and test sets, without explicit control over MLSL or SLSL. Consequently, the random split yields the highest number of interactions involving either shared enzymes or shared small molecules between the training and test sets. If shared substructures (e.g., a benzene ring within a family of molecules) were also considered, this number increases even further; however, we report only the number of exact shared interaction partners. In the enzyme-based split, the size of SLSL interactions (66,927) is twice that of MLSL interactions (26,943) for small molecules. In contrast, in the small molecule-based split, the size of SLSL interactions (13,985) is approximately seven times smaller than MLSL interactions (76,922) for enzymes. The label-based split is designed to minimize the similarity leakage within each label set (see Section 4.1.5 for details). In this split, the MLSL is smaller than SLSL for both enzymes and small molecules; however, the number of interactions involving shared enzymes or small molecules is reduced by half compared to the random split. In contrast, in the two-dimensional split, there is no MLSL and SLSL at either level between the training and test sets.

**Table 3.** Number interactions with a shared molecule/enzyme between training and test sets across split methods. MLSL: Multi-label inter-sample similarity leakage. SLSL: Single-label inter-sample similarity leakage.

| Split Method | With shared enzymes |  | With shared molecules |  | Total interactions |
| --- | --- | --- | --- | --- | --- |
|  | MLSL | SLSL | MLSL | SLSL |  |
| Random | 93,097 | 18,013 | 28,680 | 72,878 | 212,668 |
| Enzyme-based | 0 | 0 | 26,943 | 66,927 | 93,870 |
| Small molecule-based | 76,922 | 13,985 | 0 | 19 | 90,926 |
| Label-based | 3,424 | 29,128 | 21,998 | 56,572 | 111,122 |
| Two-dimensional | 0 | 0 | 0 | 0 | 0 |

These observations on inter-sample similarity leakage in different splits can be explained by an analysis of betweenness centrality (BC) in the corresponding enzyme-small molecule networks (Figure 3). BC in the training sets shows a roughly similar distribution across all split regimes. In contrast, BC has different distributions across all split regimes in the test sets (Figure 3). Most small molecules (at least 50%) in the training and test sets have a low BC and thus are structurally unimportant in terms of network connectivity, as the first quartile is equal to the median. The distribution of BC in the upper quartile changes with the split method for both enzymes and small molecules in the test sets. Enzymes generally have higher BC values than small molecules in all training sets, and the same is true for the test sets; however, the distributions are narrower, suggesting fewer extreme central nodes. Across the test sets, the median BC of small molecules remains constant, while the median BC of enzymes varies for different split methods (see additional analysis using node degree for each split method in the supplementary material figure 1).

**Fig. 3.**
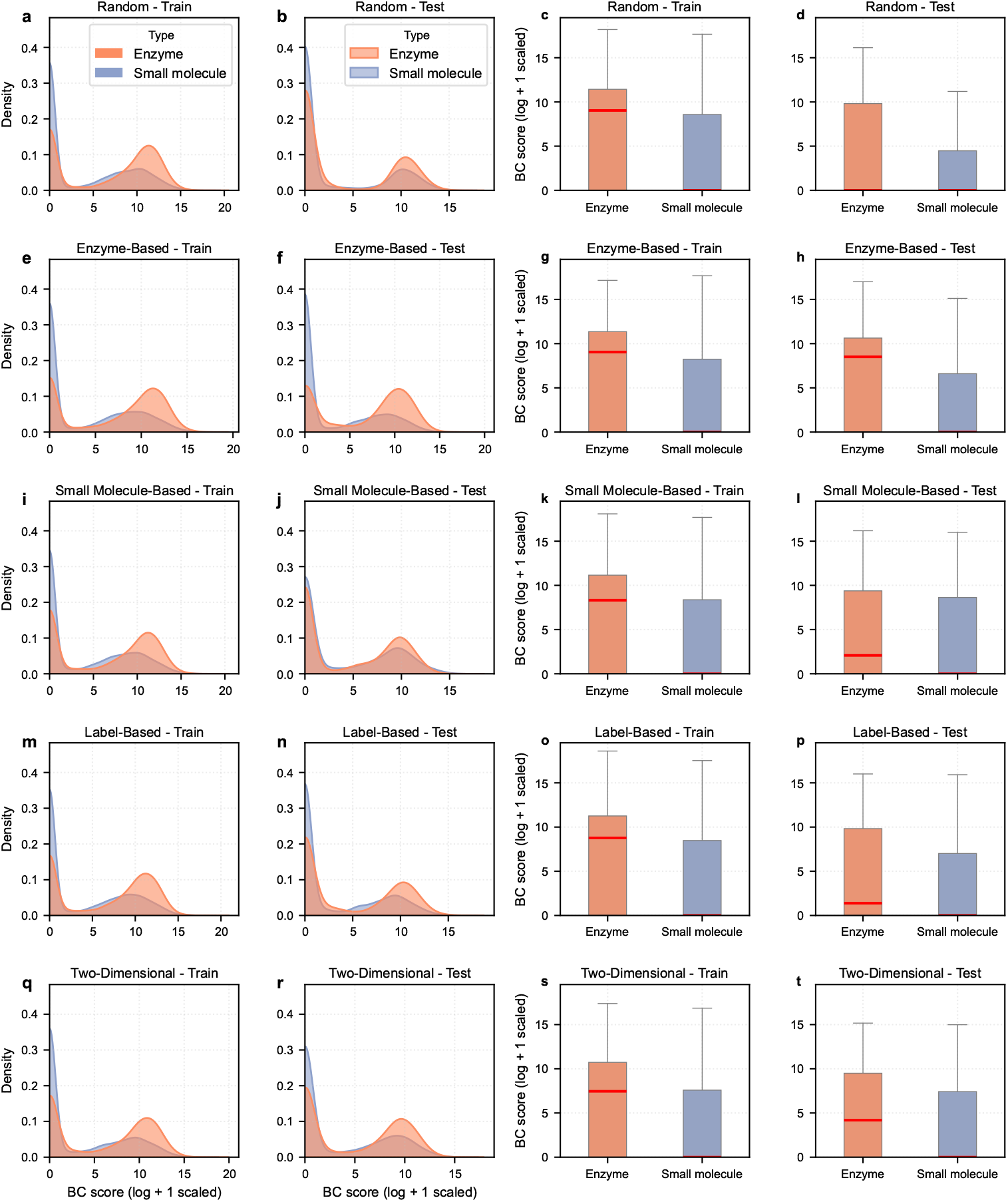
Betweenness centrality distributions for each split method. The red line indicates the median, and the whiskers represent the range of non-outlier values in the box plot

### 2.4 Model performance

The EMMA model is a dual-stream transformer with separate streams for enzyme and small molecule representations. It respects the classical transformer block structure (attention → residual connection and normalization → feed-forward → residual connection and normalization), but replaces the standard self-attention with a composite of self- and cross-attention mechanisms within each stream. The two streams are connected through a bidirectional cross-attention module, where each stream attends to the other (enzymes attend to molecules and molecules attend to enzymes) to capture mutual interaction signals. This dual-stream transformer is followed by two task-specific prediction heads: one for predicting the existence of an interaction (interaction task) and one for identifying the type of interaction (substrate or inhibitor; subclass task), both trained with focal loss functions (see Figure 7 and Section 4.3 for further details).

#### 2.4.1 Benchmarking EMMA against a random forest baseline

We trained EMMA and a random forest (RF) baseline on the five splits described above. The performance of the models varies depending on whether similarity leakage is controlled at the enzyme level, the small-molecule level, or both, or not at all, as in the random split (Table 4). This is in accordance with our earlier work, where we demonstrated that the amount of data leakage is directly correlated with performance metrics such as AUROC and accuracy ([23]).

**Table 4.** Comparison of EMMA and the Random Forest (RF) across different data split methods. We report performance on the interaction task (IT) and subclass task (ST). Results are averaged over five runs with different random seeds, with standard deviation shown. The best-performing model for each split and task is highlighted in **bold. Note:** We trained two separate random forest classifiers, one for each task. AUROC: Area under the receiver operating characteristic curve

| Split Method | Model | Task | AUROC (↑) | Accuracy (↑) | F1 score (↑) |
| --- | --- | --- | --- | --- | --- |
| Random | RF | IT | **$0.976 \pm 0.000$** | **$0.915 \pm 0.001$** | **$0.910 \pm 0.001$** |
|  |  | ST | **$0.989 \pm 0.000$** | **$0.957 \pm 0.001$** | **$0.958 \pm 0.001$** |
| | EMMA | IT | $0.944 \pm 0.001$ | $0.859 \pm 0.001$ | $0.845 \pm 0.001$ |
| | | ST | $0.984 \pm 0.000$ | $0.947 \pm 0.001$ | $0.948 \pm 0.002$ |
| Enzyme-based | RF | IT | **$0.954 \pm 0.000$** | **$0.878 \pm 0.002$** | **$0.872 \pm 0.002$** |
| | | ST | $0.980 \pm 0.000$ | **$0.937 \pm 0.001$** | **$0.940 \pm 0.001$** |
| | EMMA | IT | $0.917 \pm 0.002$ | $0.821 \pm 0.003$ | $0.810 \pm 0.004$ |
| | | ST | **$0.991 \pm 0.001$** | $0.934 \pm 0.005$ | $0.940 \pm 0.005$ |
| Small molecule-based | RF | IT | $0.734 \pm 0.017$ | $0.633 \pm 0.016$ | $0.681 \pm 0.012$ |
| | | ST | $0.889 \pm 0.003$ | $0.796 \pm 0.011$ | $0.814 \pm 0.011$ |
|  | EMMA | IT | **$0.854 \pm 0.005$** | **$0.755 \pm 0.010$** | **$0.772 \pm 0.004$** |
|  |  | ST | **$0.947 \pm 0.006$** | **$0.846 \pm 0.011$** | **$0.878 \pm 0.007$** |
| Label-based | RF | IT | $0.836 \pm 0.007$ | $0.749 \pm 0.005$ | $0.714 \pm 0.004$ |
| | | ST | $0.849 \pm 0.005$ | $0.591 \pm 0.006$ | $0.363 \pm 0.015$ |
|  | EMMA | IT | **$0.845 \pm 0.006$** | **$0.753 \pm 0.006$** | **$0.726 \pm 0.008$** |
|  |  | ST | **$0.950 \pm 0.009$** | **$0.881 \pm 0.040$** | **$0.876 \pm 0.047$** |
| Two-dimensional | RF | IT | $0.727 \pm 0.012$ | $0.648 \pm 0.014$ | $0.704 \pm 0.009$ |
| | | ST | $0.908 \pm 0.003$ | $0.769 \pm 0.005$ | $0.765 \pm 0.005$ |
|  | EMMA | IT | **$0.847 \pm 0.008$** | **$0.760 \pm 0.008$** | **$0.768 \pm 0.017$** |
|  |  | ST | **$0.954 \pm 0.004$** | **$0.884 \pm 0.011$** | **$0.899 \pm 0.012$** |

Interestingly, the RF model performs better than EMMA for a random split for both the interaction and subclass tasks, with an area under the receiver operating characteristic curve (AUROC) scores above 0.97 for the interaction task (IT) and subclass task (ST) (Table 4). However, random splits are known to inflate performance estimates by allowing highly similar molecules or enzymes to appear in both training and test sets, causing data leakage [23]. Consistent with this, we observed the highest degree of similarity leakage under the random split (Figure 4).

**Fig. 4.**
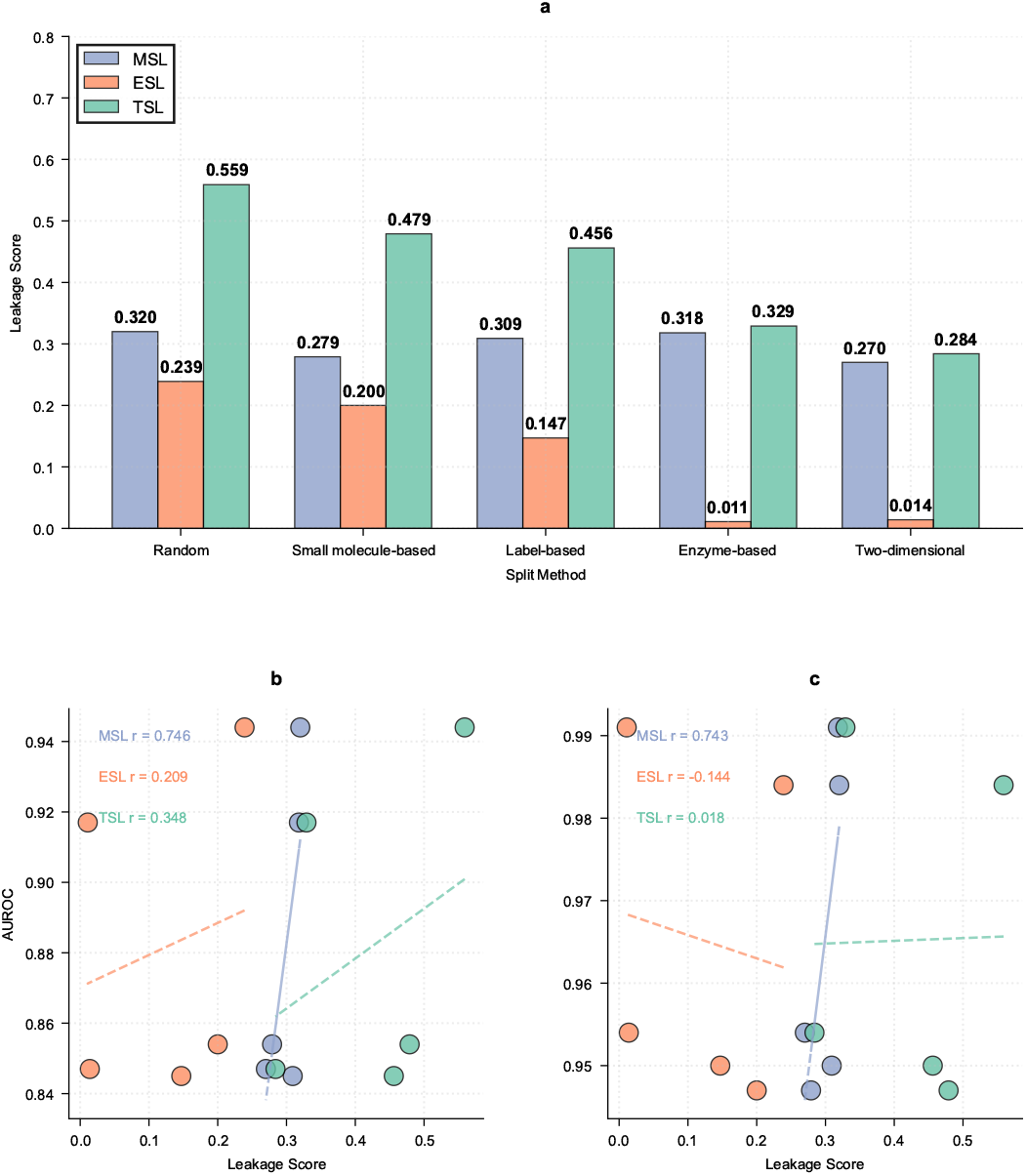
(**a**) Train-test similarity-based data leakage for each splitting method. (**b**) Pearson correlation between the AUROC score and data leakage for the interaction prediction head. (**c**) Pearson correlation between the AUROC score and data leakage for the subclass head. MSL: small molecule-similarity leakage, ESL: enzyme-similarity leakage.

In the enzyme-based split, the RF retains a modest advantage for interaction prediction, but EMMA surpasses it in the subclass task, achieving the highest AUROC (0.991). This indicates that EMMA captures more generalizable patterns when unseen enzymes are introduced. The gap becomes more pronounced in the small molecule-based split, where the RF performance degrades substantially (AUROC 0.734 for IT, AUROC 0.889 for ST), while EMMA maintains high scores (AUROC: 0.854 for IT, AUROC: 0.947 for ST). This demonstrates that EMMA generalizes more efficiently when challenged with unseen chemical scaffolds.

The label-based and two-dimensional splits represent the most stringent evaluation settings. In the label-based split, the RF performance further drops in both tasks (AUROC: 0.836 IT, AUROC: 0.849 for ST), whereas EMMA remains robust (AUROC: 0.845 for IT, AUROC: 0.950 for ST). A similar pattern is observed for the two-dimensional split, where EMMA consistently outperforms RF across both tasks. These results highlight EMMA’s ability to capture transferable representations that support generalization beyond the training distribution. These results further support our assumption that for predicting enzyme—ligand interactions, tree-based models are more sensitive to sample-similarity leakage than neural network-based models. This is due to the inherent imbalance in network node connectivity and the tendency of tree-based models to prioritize features associated with highly connected nodes. Such features frequently appear in decision splits that maximize the reduction of the splitting criterion, leading to higher feature importance.

#### 2.4.2 Effect of split regime characteristics on EMMAs performance

Our results highlight the importance of different forms of inter-sample similarity leakage in enzyme—small molecule interaction prediction. It seems that learning transferable representations for unseen small molecules is considerably more challenging than for unseen enzymes. However, measuring data leakage (Equation (3)) reveals that the total similarity leakage (TSL) in the small molecule-based split is higher than for the enzyme-based split (0.32 vs. 0.479, Figure 4). An explanation for these observations can be found in the ratio of enzymes to small molecules in the EMMI dataset (Table 1), which is 0.54, creating inherent challenges for splitting the data. As mentioned above, the enzyme—small molecules interaction datasets can be represented as a bipartite graph (Section 4.1.6). When applying small molecule-based splitting to this structure, the high connectivity of enzymes across small molecule partners frequently creates bridges between training and test partitions, as many enzymes interact with multiple small molecules. Their inclusion in training and test sets through different small molecule partners introduces data leakage, which cannot be avoided when the splitting procedure is based on small molecules. This architectural constraint explains why small molecule-based splits consistently show higher data leakage than enzyme-based splits in our experiments.

Despite the high leakage observed in the small molecule–based split, why does the model trained on the enzyme-based split achieve higher performance? The key limitation of the small molecule–based split is that it allows inter-sample similarity leakage at the enzyme level. Additionally, because the number of multi-label enzymes is larger than the number of multi-label small molecules (Figure 1), the size of the MLSL is greater than the SLSL for enzymes in the small molecule–based split compared to the enzyme-based split. As discussed in section 2.2, MLSL can negatively affect model performance when the nodes are highly connected (e.g., have a high betweenness centrality (BC)). The model performance decreases for nodes with very low (bins 1–3) or very high (bins 7–10) BC (as shown Figure 5), particularly for those splits designed to minimize similarity leakage, while the highest performance is observed for nodes with moderate BC (bins 4–6) across all splits. Our results confirm that as the BC of enzymes increases, model performance decreases. Since most highly connected enzymes tend to have multiple labels (Figure 1a), we conclude that the larger MLSL for enzymes in the small molecule–based split is responsible for its lower performance compared to the enzyme-based split.

**Fig. 5.**
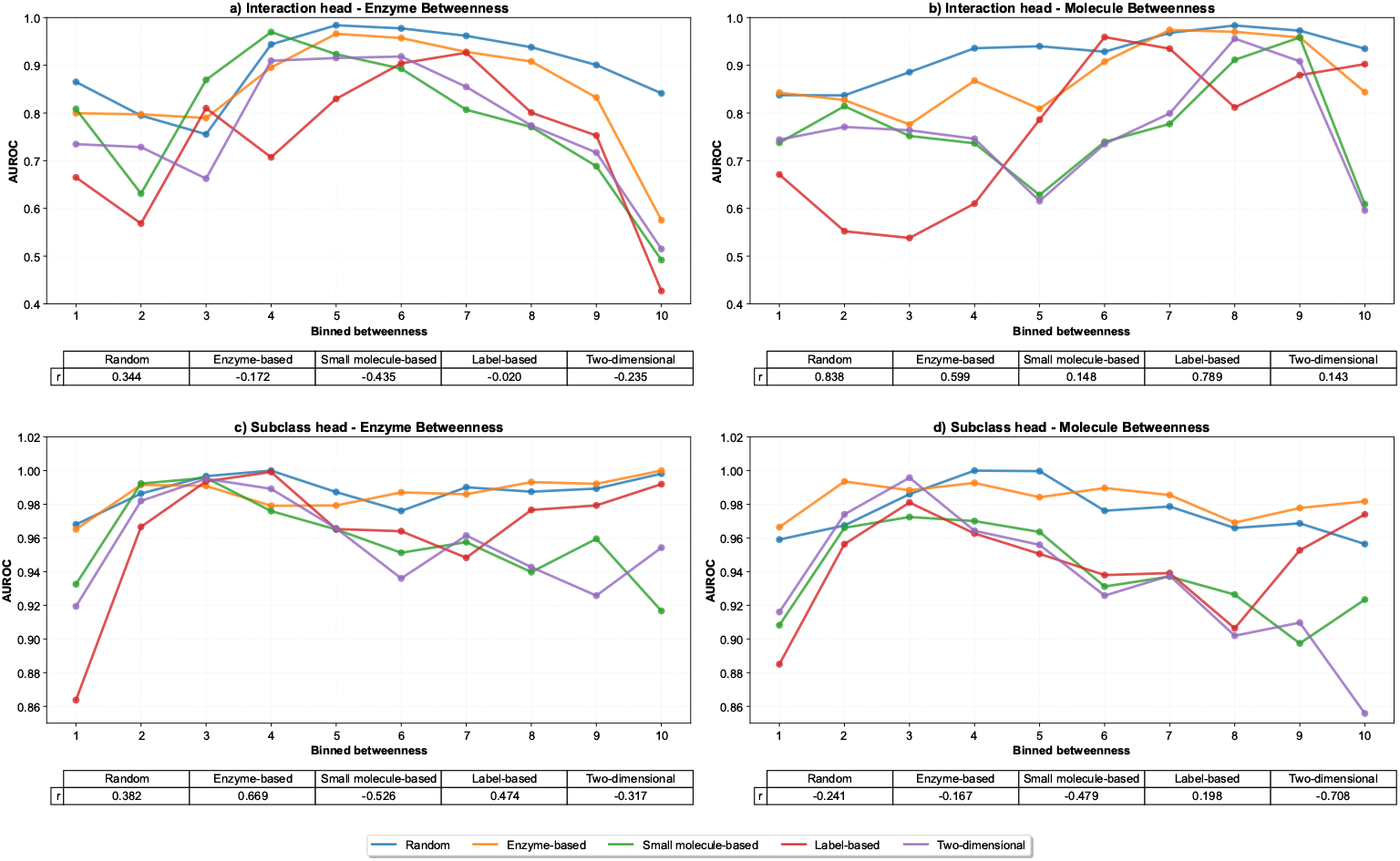
Model performance (AUROC) across different levels of betweenness centrality. Nodes were grouped into bins according to their log-transformed betweenness centrality, and AUROC was computed within each bin for both enzyme- and molecule-level betweenness. Results are shown separately for the interaction and subclass heads across all data splits. Tables beneath each subplot report the Pearson correlation coefficients (*r*) between AUROC and betweenness bin indices for each split.

Interestingly, in the label-based split, where multi-label enzymes and small molecules can appear in both the training and test sets while minimizing similarity leakage within each label set, the model achieved lower performance compared to the enzyme- and small molecule-based splits, despite showing an intermediate TSL of 0.456. The size of MLSL is smaller than SLSL at both the enzyme and small molecule levels for this split (Table 3). Here, the enzyme–substrate set was divided using a small molecule-based split because its enzyme-to-small molecule ratio is greater than 1, indicating that small molecules are less connected on average; thus, splitting by small molecules better minimizes inter-sample similarity leakage. Conversely, the enzyme–inhibitor and enzyme–non-interacting sets were divided using an enzyme-based split, as their ratios are less than 1 and enzymes are more closely connected, making enzyme-based splitting more effective at reducing leakage. By splitting the data into these three smaller problems rather than splitting the entire dataset at once, the leakage within each subset was effectively minimized. However, leakage across label sets still occurs in the form of MLSL at both enzyme and small molecule levels.

In the two-dimensional split, the most challenging evaluation scenario, which minimizes similarity leakage across both enzymes and small molecules simultaneously, the model achieves a comparably low performance, reflecting the inherent difficulty of predicting interactions for entirely novel enzyme—small molecule combinations. In this split, the distributions of BC for both the training and test sets are narrower, suggesting fewer extreme central enzymes and small molecules compared to the one-dimensional splits. This stems from the fact that the two-dimensional split removes interactions that bridge between training molecules and test molecules, to minimize inter-sample similarity leakage. In our analysis, the number of multi-label enzymes decreased by 2% in the combined train–test data of the two-dimensional split compared to the EMMI dataset. In contrast, the number of multi-label small molecules showed only a slight decrease.

Importantly, the ST consistently achieves better performance metrics than the IT across all splitting strategies. This difference may be rooted in how the dataset was constructed: interacting pairs (substrates and inhibitors) are supported by strong experimental evidence, either through direct biochemical evidence or experimentally validated functional GO annotations (see Section 4.1). By contrast, enzyme—non-interacting pairs are defined indirectly, via taking pairs with low binding affinity in experimental assays. As a result, the likelihood of mislabeling is higher for enzyme—non-interacting pairs than for interacting ones. Consequently, subclass-level classification benefits from more reliable ground-truth labels, whereas binary interaction tasks suffer from noise introduced by uncertain non-interacting assignments. As systematic data on enzyme—non-interacting pairs is lacking, we believe that our strategy for generating negative data—incorporating experimentally validated enzyme—non-interacting ligand pairs—has borne fruit across different split regimes. This emphasizes the value of integrating experimental evidence and reducing reliance on synthetic negatives to ensure biologically meaningful predictions. Furthermore, the higher number of multi-label nodes in the interaction dataset, compared to the sub-class dataset (Table 1, makes the interaction task inherently more difficult than the subclass task.

## 3 Discussion

In this study, we introduce EMMA, a Transformer-based multi-task learning framework that leverages a novel task formulation to jointly predict two distinct yet related binary tasks within a shared latent space using two classification heads: (i) an interaction head that distinguishes low-affinity pairs from high-affinity ones, and (ii) a subclass head that further classifies high-affinity interactions as either inhibitory or catalytic. To train and validate the model, we compiled a comprehensive dataset that incorporates experimentally validated enzyme—small molecule pairs not only for substrate and inhibitor pairs, but also for non-interacting ones. This biologically grounded approach yielded exciting results under different split regimes, with EMMA achieving an average AUROC of 0.866 for the interaction task and 0.960 for the subclass task, demonstrating a high ability at rank prediction.

We specifically studied the effect of inter-sample similarity leakage on model performance and showed that higher leakage does not always improve performance. We further divided the leakage into single-label and multi-label inter-sample similarity leakage (SLSL and MLSL, respectively). We demonstrated how SLSL can artificially enhance model performance, whereas MLSL is harmful for it. Furthermore, we demonstrated that the topology of the enzyme—small molecule interaction network, represented as a bipartite graph, and how it is separated into the training and test splits, could amplify or reduce the effect of the MLSL and SLSL. In particular, nodes with high connectivity are prone to be memorized, as when similarity leakage is minimized, performance on nodes with high connectivity—often corresponding to multi-label nodes—declines markedly.

We believe that the model’s performance on the single-and multi-label nodes with different degrees of BC can provide insight into its ability to learn and generalize. To the best of our knowledge, there is currently no established metric to evaluate this aspect of model capability. We believe our findings offer a foundation for developing such a metric. Our analysis emphasizes the complexity of predicting enzyme—substrate and enzyme—inhibitor interactions, where special attention needs to be paid to training regimes, interaction network topology, and data splitting to ensure models’ generalizability to unseen enzymes and small molecules.

## 4 Methods

### 4.1 Creating the EMMI dataset

First, we have compiled a dedicated dataset of Enzyme-sMall Molecule Interactions (EMMI). The EMMI dataset contains experimentally confirmed enzyme—substrate, enzyme—non-interacting, and enzyme—inhibitor pairs. To create this dataset, we extracted data from the following sources: BindingDB [29], Brenda [30], ChEMBL [31, 32], Gene Ontology [33, 34], Iupher-PBS [35], PubChem [36], Rhea [37], Sabio-RK [38], and UniProt [39] (Figure 6).

**Fig. 6.**
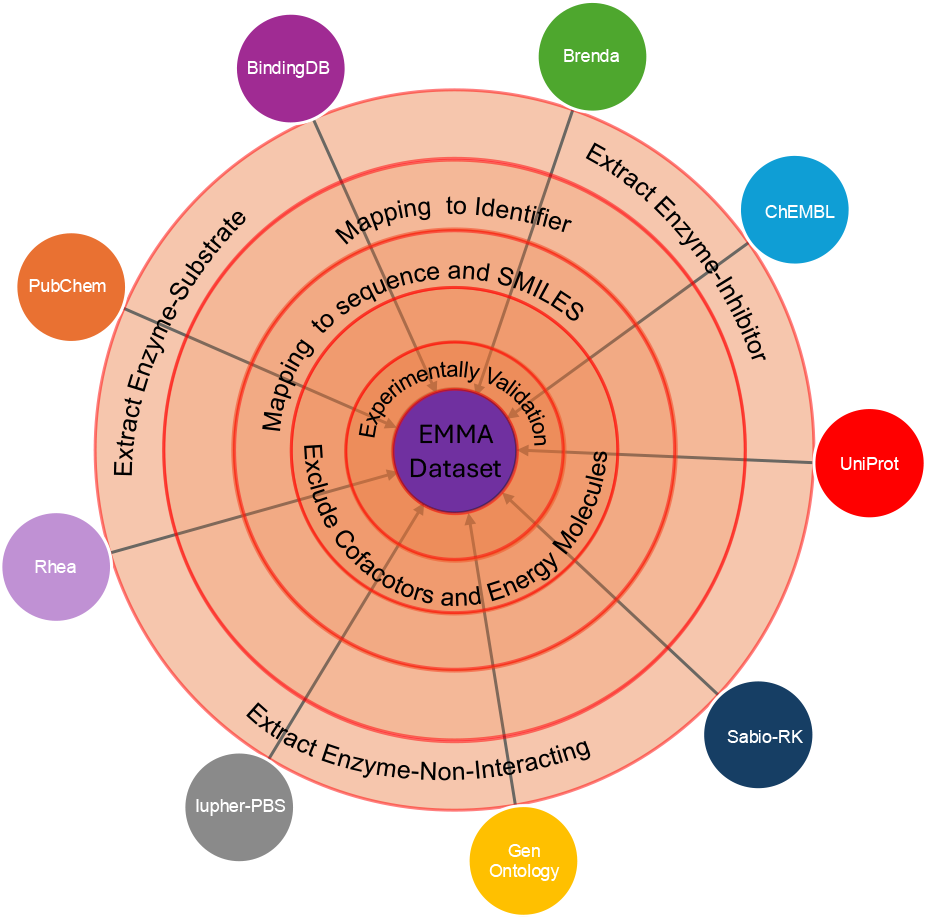
Abstract view of data processing steps to create EMMI dataset from various sources.

In the data-processing pipeline, first, all enzyme–small molecule pairs were extracted and each entity was mapped to a unique identifier. For enzymes, UniProt IDs were provided by the source databases. For small molecules, however, most of the sources provided only an IUPAC name or a common name. To map small molecule names to standardized identifiers, if the source database did not provide an identifier, we first attempted to retrieve PubChem Compound IDs (CIDs) using the PubChem PUG REST API [40]. For small molecule names that lacked a CID, we created a manually curated mapping. This dictionary integrates data from several databases (ChEBI ontology [41], HMDB [42], SMPDB [43], and KEGG [44]), and contains 610,608 unique small molecule names mapped to IDs. Our mapping strategy prioritizes a PubChem ID; if none is found, we then search for a ChEBI ID, followed by a KEGG ID.

These identifiers were then used to associate enzymes with sequences and small molecules with SMILES strings. Only pairs with experimentally validated annotations were retained (see Section 4.1.1 for details). Pairs with invalid sequences or SMILES strings were excluded. Since the maximum position embeddings (also called token size limit) in ESM2 t30 [45] and MolFormer XL [46] are 1,024 and 202, respectively, we only retained enzyme sequences of up to 1,500 amino acids ( ~ 47% longer than the embedding limit of the ESM2 t30), with only ~ 11% of enzymes in the EMMI dataset have a length between 1,025 and 1,500 amino acids. For SMILES strings, although the majority already fall well below the 202-token limit, we permitted SMILES strings of up to 512 characters; as a result, only less than 1% of small molecules exceed the token limit in the EMMI dataset. These thresholds were chosen to minimize the number of truncated enzymes or small molecules while maximizing the number of enzymes, thereby achieving a more balanced enzyme-to-molecule ratio within each label set (e.g., enzyme—substrate, enzyme—inhibitor, and enzyme—non-interacting) and in the EMMI dataset. SMILES strings were subsequently canonicalized using RdKit [47], and duplicates were removed at the level of enzyme–small molecule pairs. Finally, we filtered out all interactions involving known coenzymes, cofactors, and energy-transfer small molecules (Supplementary table 1). Details of the data mining and processing procedures for each database are provided in the Supplementary material Section S1.

#### 4.1.1 Experimentally confirmed data points

We distinguish between two types of enzyme—small molecule pairs. The first category includes enzyme—small molecule pairs supported by quantitative binding or functional assay measurements, referred to as binding affinity-related assay (BAA) pairs. For those interactions, one or multiple of the metrics in Table 5 have been measured. These assays provide direct evidence of molecular interaction. We refer to these pairs as BAA-confirmed pairs.

**Table 5.** Descriptions of the experimental measurements for enzyme-small molecules interactions.

| Name | Description |
| --- | --- |
| $K_i$ | The *inhibition constant* is a direct measure of inhibitor binding affinity, independent of assay conditions. |
| $K_d$ | The *dissociation constant* indicates ligand binding strength (e.g., from surface plasmon resonance (SPR) or isothermal titration calorimetry (ITC)). |
| $K_a$ | The *association constant* is the inverse of $K_d$, quantifying binding affinity. |
| $K_m$ | The *Michaelis constant* reflects enzyme-substrate affinity in catalytic reactions. |
| $IC_{50}$ | The *half-maximal inhibitory concentration* is a functional measure of inhibition, dependent on assay setup (e.g., substrate concentration, incubation time). |
| $EC_X$ | The *X-Maximal effective concentration* is a functional measure of the concentration of a compound that produces X% of its maximal biological effect. Like $IC_{50}$, it is dependent on assay conditions and reflects apparent potency rather than an intrinsic binding constant. |
| **Inhibition** | Reports the extent of enzyme activity reduction either as a percentage at a given concentration, or as the compound concentration required to achieve a defined level of inhibition. |
| **MIC** | The *Minimum Inhibitory Concentration* defines the lowest concentration of a compound required to prevent visible growth of a microorganism. |

**Table 6.**
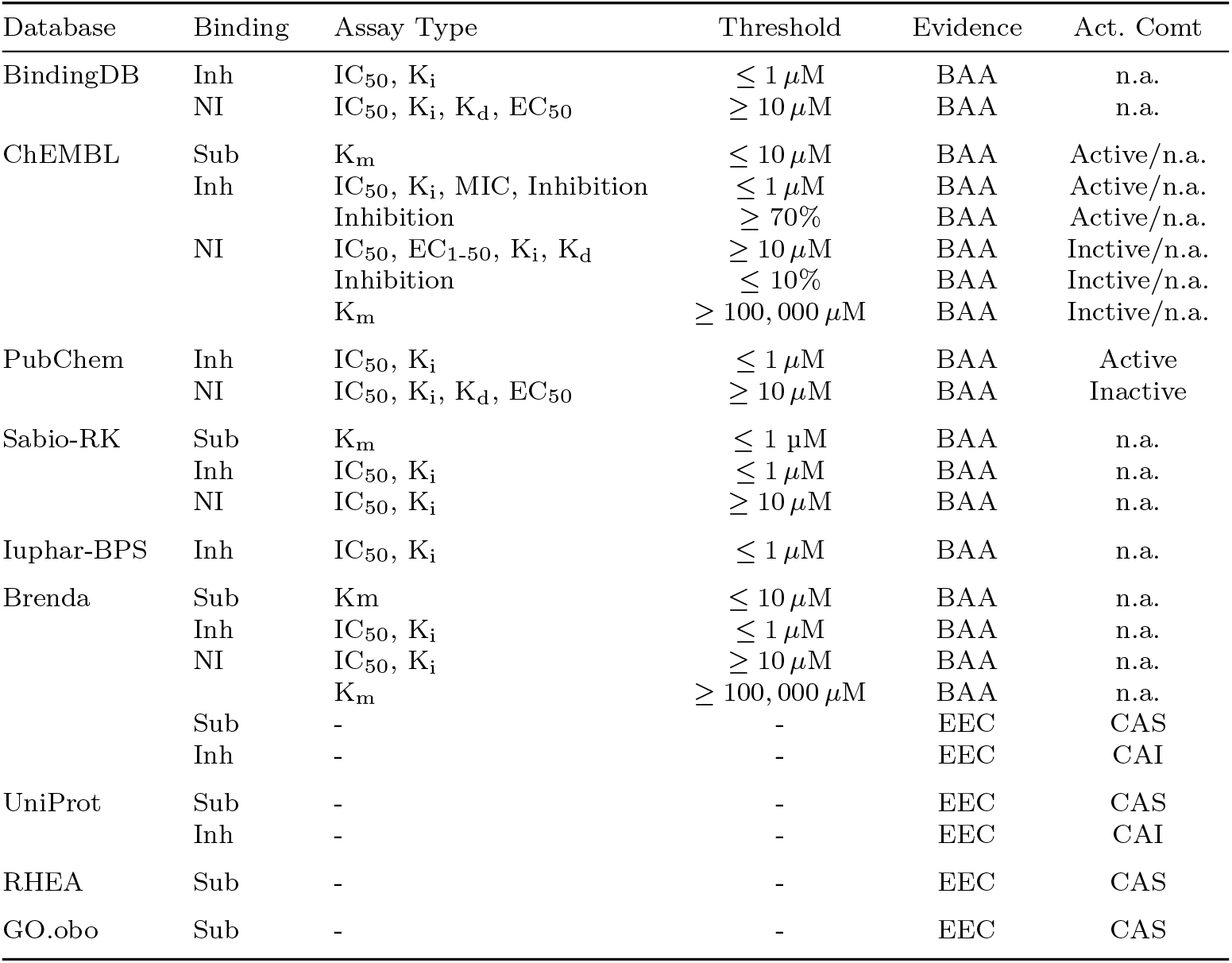
Database filtering criteria with assay type specifications. **Note:** All IC_50_ values were converted to K_i_. To indicate the source of each measurement, the original assay names from the respective databases were retained. Inh: inhibitor; Sub: substrate; NI: non-interacting; BAA: binding affinity-related assay; EEC: experimental evidence code; Act. Comt: activity comment; CAS: classified as a substrate; CAI: classified as an inhibitor; n.a.: not available.

The second category consists of enzyme–small molecule pairs inferred from experimentally validated Gene Ontology annotations (GOA). In this case, the presence of a functional GO ID with associated *experimental evidence code* (EEC) for an enzyme implies functional relevance of the corresponding interacting small molecule. To map functional GO IDs to EECs, we processed the ~ 1.2 billion GOAs and selected only those supported by EECs. This curated dataset was then used as a filter to retain only enzyme–small molecule pairs for which both the UniProt ID and the functional GO ID could be jointly mapped to an EEC-supported term. We refer to them as EEC-confirmed pairs.

This classification framework allows us to capture both direct biochemical evidence and biologically supported associations.

#### 4.1.2 Conversion of experimentally measured values

The conversion of IC_50_ to K_i_ is necessary because K_i_ is a thermodynamic constant that reflects the true binding affinity of an inhibitor, independent of assay conditions such as substrate concentration or reaction time. In contrast, IC_50_ values are assay-specific and cannot be directly compared across different studies without normalization.

According to Kalliokoski et al. [48], although IC_50_ values are more abundant in public datasets like ChEMBL, they are only moderately more variable than K_i_ values (by approximately 25%). The authors showed that IC_50_ values from mixed assays can be used reliably in large-scale analysis if converted appropriately, and that the average ratio between IC_50_ and K_i_ values across many protein–ligand pairs is approximately 2, which corresponds to assay conditions where the substarte concentration [*S*] = K_m_ in the Cheng-Prusoff equation [49]:

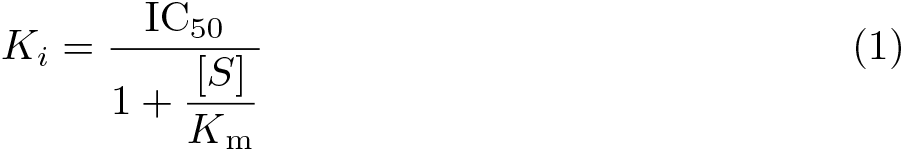

Thus, converting IC_50_ to K_i_ improves comparability of inhibitor-binding data in our integrated dataset. This heuristic approximation also avoids explicit dependence on substrate concentration or K_m_ especially for working with large-scale heterogeneous databases (e.g., PubChem, ChEMBL), where assay-specific details such as [S] and K_m_ are often missing or unstandardized. While it is an approximation, it preserves the relative ranking of binding affinities across different inhibition levels and enables effective comparison of IC_50_ and K_i_ values within a unified framework.

We also included direct conversions of K_a_ to K_d_ using:

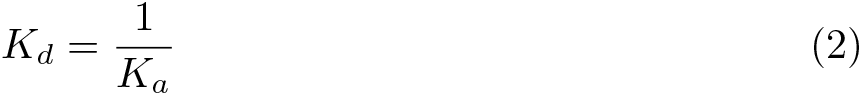

For EC_X_ assays, we did not find a standard or reliable method for converting values to K_i_ or K_d_. Therefore, we retained the originally reported values. Furthermore, EC_X_ values can reflect the effective concentration required to elicit a biological response, either from an inhibitor or a substrate. For large-scale analysis, it is not feasible to determine from the assay descriptions whether the reported EC_50_ value is for an inhibitory or catalytic interaction. Therefore, we used them only to identify noninteracting pairs and considered only EC_X_ values where 1 ≤ *X* ≤ 50.

#### 4.1.3 Assigning active and inactive interaction

For BAA-confirmed pairs, activity comments, when available, are used to label pairs as active or inactive based on the assay values. In PubChem, a general threshold of 10 µM is applied for BAAs (except for K_m_) where pairs with values above 10 µM are considered as inactive, while values below this threshold indicate interaction [50]. However, exceptions exist; in some cases, some pairs from functional assays with assay values above 10 µM are annotated as active, while others with values below 10 µM are labeled as inactive. These exceptions are assay-specific and comparable only under certain conditions.

These exceptional cases make it risky to assign labels based on a universal threshold, especially for data points without activity comments. However, despite these limitations and the assay-specific nature of functional measurements, lower assay values generally indicate a stronger binding affinity and the higher values indicate a weaker binding affinity for BAAs. Hence, we defined a number of binding strength categories as follows: strong (≤ 1 *µ*M), moderate to weak (1 *µ*M *< x <* 10 *µ*M), and very weak ≥ 10 *µ*M, corresponding to BAA, except K_m_. We only selected pairs with strong and very weak binding affinity for the EMMI dataset: pairs with assay values ≥ 10 *µ*M are labeled as non-interacting, while pairs with values ≤ 1 *µ*M are labeled as interacting.

The conversion of IC_50_ to K_i_ values could further effectively minimize the risk for IC_50_ assay above 10 µM which are annotated as active (Section 4.1.2). Furthermore, for functional assays with values below 10 *µ*M that are labeled as inactive, the 1 *µ*M cutoff is sufficiently low to minimize the number of such assays that could be incorrectly classified as interacting.

The K_m_ values of enzymes vary widely. For most enzymes, K_m_ lies between 0.1 µM and 100,000 µM. The median K_m_ is approximately 100 µM, with about 60% of all K_m_ values falling in the range of 10–1000 µM [51, 52]. To define high-affinity and moderate interactions, we selected pairs with K_m_ ≤ 10 µM. To mitigate potential noise, we define pairs with K_m_ ≥ 100, 000 µM as non-interacting pairs. The thresholds for the MIC and inhibition assays were derived through an empirical assessment of the ChEMBL database, based on assays for which activity comments are available.

Furthermore, within the interacting pairs, for those pairs with a K_i_, IC_50_, MIC or inhibition value or classified as inhibitor (CAI), we sub-labeled as enzyme—inhibitor pairs. Similarly, enzyme—small molecule pairs with a K_m_ value or classified as a substrate (CAS) are sub-labeled as enzyme—substrate.

#### 4.1.4 Clustering-based down-sampling

After data labeling, to ensure balanced learning, we implemented clustering-based down-sampling for enzymes with a high number of associated inhibitors and non-interacting small molecules (Table 1). We first converted all small molecules to extended-connectivity fingerprints (ECFPs). Specifically, we computed Morgan fingerprints with a radius of 3 (ECFP6), a bit length of 2048, and chirality enabled using RdKit [47]. For each enzyme, the associated small molecules were clustered based on their fingerprints using the *K*-means clustering algorithm, with a fixed random seed to ensure deterministic results. The global number of clusters *K* per enzyme was tuned to balance the classes (four clusters for inhibitors and six for non-interactors). From each cluster, we selected the small molecule closest to the cluster centroid. In cases where multiple small molecules were equidistant from the centroid, a deterministic tie-breaking procedure ensured reproducibility.

This strategy allowed us to control label imbalance and retain structurally diverse representatives of both inhibitors and non-interacting small molecules for each enzyme. The composition of the EMMI dataset, including the contribution of each source database after cluster-based down-sampling, is summarized in Supplementary table 2.

#### 4.1.5 Data splitting and data leakage

We split the EMMI dataset into training (80%) and test (20%) sets using the following methods implemented in DataSAIL [23]: enzyme-based, small molecule-based splits, and a two-dimensional split (Table 2). In the enzyme-based split, highly similar enzymes are confined to either the training or test set; and MLSL and SLSL can occur at the small molecule level (Table 3). Likewise, in the small molecule-based split, structurally similar small molecules are restricted to a single set to prevent overlap; however, MLSL and SLSL can happen at the enzyme level. In the two-dimensional split, no similar enzyms and small molecules appear in more than one split, and consequently, no form of inter-sample similarity leakage (MLSL or SLSL) is permitted.

We further propose a label-based split strategy, motivated by both the bipartite graph structure of the interaction datasets (Section 4.1.6) and a divide-and-conquer principle. In the bipartite graph, the enzyme-to-small molecule ratio acts as an indicator of which node type (enzymes or small molecules) has higher average connectivity or degree. By splitting the dataset based on the more connected partner, we minimize the number of bridges between the training and test sets. This approach effectively reduces inter-sample similarity leakage compared to splitting based on node type, which has a lower average connectivity. The divide-and-conquer principle complements this approach by treating each label set (e.g., enzyme—substrate) as a separate sub-problem. By performing splits independently within these smaller subproblems based on the enzyme-to-small molecule ratio, we can effectively control inter-sample similarity leakage than by splitting the entire dataset at once. In practice, in the EMMI dataset, since the enzyme-to-small molecule ratio is greater than 1 in the enzyme— substrate set, we split it using a small molecule-based split. Conversely, since this ratio is less than 1 in the enzyme—inhibitor and enzyme—non-interacting sets, we applied enzyme-based splits (Table 1). This process yielded three separate train/test sets, which we subsequently combined into one unified train set and one unified test set. Unlike the two-dimensional split, the label-based split allows MLSL and SLSL at both enzymes and small molecules (Table 3). However, by dividing the splitting problem into label-specific subproblems, this divide-and-conquer strategy minimizes similarity leakage more effectively within each label set than other one-dimensional splits applied to the entire dataset.

Finally, a random split was used as a baseline, in which interactions were assigned randomly to the training and test sets, allowing both MLSL and SLSL across enzymes and molecules.

For all split regimes described above, 20% of the training data were further reserved as a validation set for hyperparameter tuning.

We quantify the leakage between any two splits *s* and *s*^′^, e.g., training and test, of a dataset 𝒟 as described in Joeres et al. [23] as

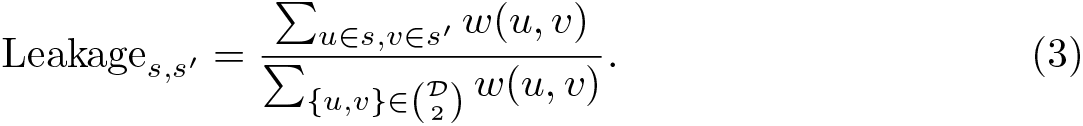

*u* and *v* are data points in 𝒟, which can be enzymes, substrates, or interactions thereof. *w* : 𝒟 × 𝒟 → ℝ is a similarity function between data points in 𝒟. The numerator measures the similarity between distinct splits, while the denominator normalizes by total similarity in the dataset. Higher leakage values indicate greater similarity between the corresponding splits.

#### 4.1.6 Analysis of the bipartite interaction graph

The enzyme—molecule interaction datasets can be formally represented as a bipartite graph where enzymes and molecules form two disjoint vertex sets, with edges representing observed interactions. We computed the exact betweenness centrality (BC) for all nodes using the Brandes algorithm and applied a log(x + 1) transformation to stabilize variance and enable comparison [53]. It allows us to quantify the extent to which individual enzymes or molecules act as bridges within the interaction landscape and how deeply the network is interconnected (Figure 3).

To further examine how nodes’ position in the interaction network of test sets affects the model performance, we conducted a binned betweenness analysis across all test sets. Nodes were sorted by their log-transformed BC and divided into equal-width bins, where higher bin indices correspond to higher average BC values. For each bin, we evaluated model performance (AUROC) separately for enzyme- and molecule-level betweenness for both the interaction and subclass tasks. This analysis enables us to assess whether predictive performance systematically varies with the structural importance of nodes within the bipartite network (Figure 5).

### 4.2 Implementation

EMMA is implemented in Python 3.12.0 [54] using PyTorch 2.6.0 [55] as the primary deep learning framework. Data processing and numerical computations rely on Pandas 2.2.3 [56] and NumPy 1.26.4 [57]. For domain-specific tasks, we employ BioPython 1.85 [58], RDKit 2025.03.2 [47], ChemDataExtractor 1.3.0 [59], ChEMBL Webresource Client 0.10.9 [60], and LibChEBIpy 1.0.10 [61]. Machine learning utilities are provided by Scikit-learn 1.6.1 [62] and SciPy 1.15.2 [63]. Visualization is performed with Matplotlib 3.10.1 [64] and Seaborn 0.13.2 [65]. Additional tools include TQDM 4.67.1 [66] for progress monitoring, DataSAIL 1.1.1 [23] for dataset splitting, Diamond 2.1.11 [67] for sequence alignment, Transformers 4.48.1 [68] for pretrained models, Fair-ESM 2.0.0 [69] for protein embeddings, and Py-cd-hit 1.1.4 [70] for sequence clustering.

Experiment tracking and logging are performed with Weights & Biases 0.19.8 [71], while Requests 2.32.3 [72] facilitates API queries, and Colorama 0.4.6 [73] enhances command-line output.

### 4.3 EMMA architecture

We developed a transformer-based multi-task learning architecture that we called the **E**nzyme–small **M**olecule interaction **M**ulti-head **A**ttention (EMMA) model, to jointly predict the strength of enzyme–small molecule interactions and classify their interaction types (Figure 7). The LXMERT encoder inspired our encoder design [74], which employs dual-stream transformer mechanisms for vision and language modalities.

**Fig. 7.**
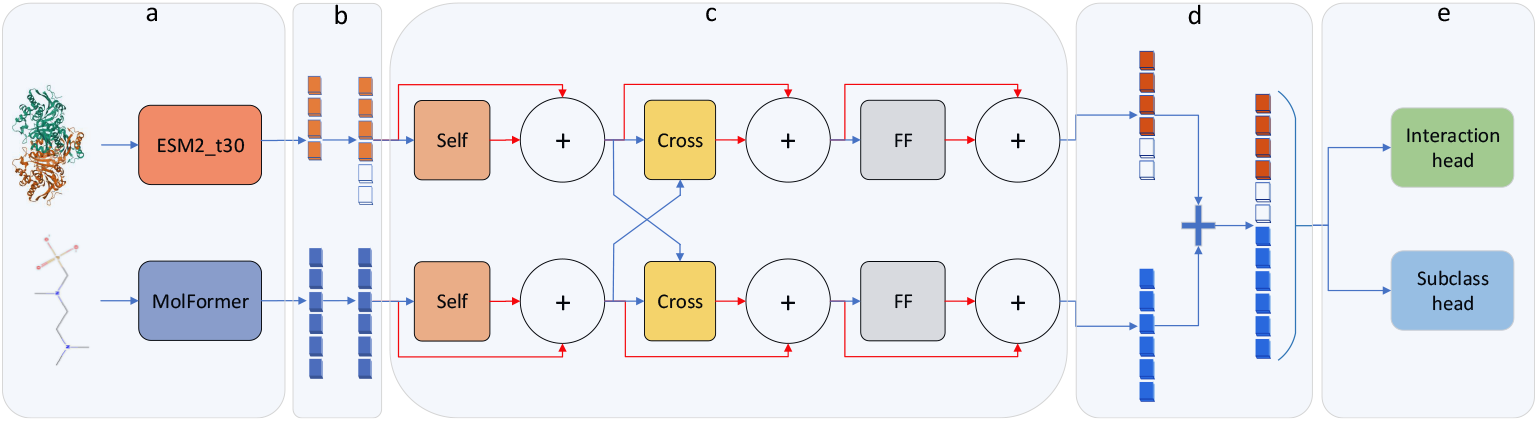
EMMA architecture. *Self, Cross*, and *FF* are abbreviations for self-attention, cross-attention, and feed-forward, respectively. Red arrows represent residual connection followed by normalization (summation sign). (**a**) Modality-specific encoders. (**b**) Projection block. (**c**) Dual-stream transformer mechanisms. (**d**) Concatenation. (**e**) Task-specific prediction heads.

Unlike LXMERT, EMMA omits intermediate feed-forward blocks inside the single-modality encoders and self-attention sub-layers inside the cross-modality encoder (see the original paper [74] for more information). We found that for the enzyme—small molecule interaction dataset, these additional layers introduced unnecessary complexity without improving performance. Instead, EMMA directly integrates the outputs of the self-modality encoders into the cross-modality encoder, simplifying the architecture while maintaining its predictive power. Our model comprises three primary components: modality-specific encoders, dual-stream transformer mechanisms, and task-specific prediction heads.

Each enzyme is represented using a single, mean-pooled, 640-dimensional embedding vector derived from the ESM2 t30 model [45], and each small molecule is encoded with a single, mean-pooled, 768-dimensional embedding vector from MolFormer XL [46] (Figure 7 a). To enable cross-attention between these two modalities, enzyme embeddings are projected into the same 768-dimensional latent space as the molecule embeddings using a learnable linear transformation, while small molecule embeddings are linearly transformed within the same dimensionality (Figure 7 b); both projections are followed by layer normalization.

To model intra-modal dependencies, self-attention layers are applied independently to enzyme and small molecule embeddings. Each stream is processed by a multihead attention mechanism with 32 heads, followed by residual connections and layer normalization. This is followed by cross-attention, where enzymes attend to the features of small molecules and vice versa, enabling the model to learn modality-specific interaction signals. Residual connections and layer normalization also follow each cross-attention. The resulting enzyme and small molecule representations are further processed through feed-forward neural networks with GELU activation and residual connections and normalization (Figure 7 c). The refined representations are then concatenated to form a 1536-dimensional interaction feature vector (Figure 7 d).

This shared representation is passed through two independent single-layer perceptrons: one for predicting binary interactions (interacting or non-interacting) and the other for subclassifying interacting pairs into enzyme-substrate and enzyme-inhibitor pairs. Each head consists of a linear layer (1024 units) followed by batch normalization, ReLU activation, dropout regularization (Figure 7 e).

The model is trained using a composite loss function. A binary focal loss [75] is applied to the interaction head, while a second focal loss is used for subclass predictions on interacting pairs only. Optimization is performed using AdamW[76] with learning rate scheduling via ReduceLROnPlateau, and gradient clipping is applied to ensure stability. To account for differences in data distribution and class balance across different dataset splits, we manually tuned key training and loss parameters for each split. Specifically, we adjusted the batch size, initial learning rate, weight decay, and the focal loss hyperparameters (gamma and alpha) for each split method. This allowed the model to better accommodate variations in sample size and label imbalance between splits, ensuring stable optimization and improved convergence for both tasks. To prevent overfitting and reduce unnecessary training time, we implemented an early stopping strategy based on the validation loss. A maximum of 100 epochs was set for all split methods. After each epoch, the model’s performance on the validation set is evaluated. If the validation loss does not improve for three consecutive epochs (patience = 3), training is terminated early.

We also implemented a RF classifier as a baseline model. Two RF classifiers were trained: one for the interaction task and one for the subclass prediction task. The models were trained with 100 trees, unlimited depth, the Gini criterion, and a minimum of two samples per split.

Evaluation metrics (AUROC, accuracy, and F1-score) were computed separately for both tasks during training and validation.

## 5 Data Availability

The train-test splits are publicly available at https://github.com/kalininalab/EMMA. EMMI dataset along with the raw and intermediate data generated during preprocessing are archived on Zenodo (https://doi.org/10.5281/zenodo.17280853).

## 6 Code Availability

All source code and implementation details are openly accessible at the GitHub repository https://github.com/kalininalab/EMMA.

## 7 Author Contribution

V.A.E. conceived the study, developed the methodology, and implemented the pipeline. O.V.K. and R.J. contributed to study conception, validation, and jointly supervised the project. All authors wrote and edited the manuscript.

## 8 Acknowledgements

R.J. was partly funded from the HelmhotzAI project XAI-Graph. O.V.K. acknowledges financial support from the Klaus Faber Foundation.

