## Supplementary material for "A General Transformer-Based Multi-Task Learning Framework for Predicting Interaction Types between Enzyme and Small Molecule"

#### Supplementary information

#### S1 Data Mining and Processing

In the following sections, we describe the data mining and processing steps for each database. When data is obtained via API access or through text mining, we flag it with 'API' or 'TM', respectively.

To map functional GO IDs to experimental evidence codes (EECs), we processed ∼1.2 billion gene ontology annotations (GOAs) and selected only those supported by EEC. From each experimentally supported GOA, we create a data frame containing the associated UniProt ID and its EEC, which we refer to as *GoToEec* (see Section the Methods for more details).

To map small molecule names to IDs using the PubChem API, we utilize our manually curated mapping dictionary. This dictionary integrates data from several databases (ChEBI ontology, HMDB, SMPDB, and KEGG) which we refer to as *MolToId* (see Section methods for more details).

##### S1.1 BAA-Confirmed Pairs

The first category includes databases that provide enzyme–small molecule pairs with associated binding affinity-related assay (BAA) measurements, such as $IC_{50}$, $EC_X$, $K_a$, $K_d$, $K_i$, $K_m$, MIC, and inhibition values. We refer to them as *BAA-confirmed pairs.* Data of this type were extracted from BindingDB, Brenda, ChEMBL, IUPHAR-PBS, PubChem, and Sabio-RK.

###### S1.1.1 BindingDB Database

- **Extraction Type:** TM.
- **Data Mining:** BindingDB provides data files in a tab-separated value (TSV) format, where each row contains information for one binding measurement. For each binding measurement, the extracted information includes:
  - **UniProt ID**: A unique identifier for each protein.
  - **PubChem CID**: PubChem CID for ligand.

  - **SMILES string**: SMILES string for ligand.
  - **Measured affinity**: $K_i$, $IC_{50}$, $K_d$, and $EC_{50}$.

- **Data Mapping:**
  - UniProt ID → enzyme commission (EC) number via UniProt API.
  - UniProt ID → sequence via UniProt API.

- **Data Filteration:**
  - Retain only interactions with an EC number and a measured affinity.
  - Exclude cofactor and energy-transfer small molecules.

- **Final curated dataset:** It contains 1,723,862 enzyme-small molecule pairs.

##### S1.1.2 Brenda Database

- **Extraction Type:** TM.
- **Data Mining:** In the first stage, we extracted the following information for each catalytic activity.
  - **UniProt ID**
  - **EC number**
  - **Substrate name**
  - **Inhibitor name**
  - **Measured affinity**: $K_i$, $IC_{50}$, and $K_m$.

- **Data Mapping:**
  - UniProt ID → sequence via UniProt API.
  - Substrate/Inhibitor name → an ID via PubChem API and MolToId.
  - Substrate/Inhibitor ID → SMILES string (via PubChem, ChEBI, or KEGG APIs)

- **Data Filteration:**
  - Retain only interactions with an EC number and a measured affinity.
  - Exclude cofactor and energy-transfer small molecules.

- **Final curated dataset:** It contains 16,480 enzyme-small molecule pairs.

##### S1.1.3 ChEMBL Database

- **Extraction Type:** API.
- **Data Mining:** To extract high-confidence data from ChEMBL, we connected to various clients in the ChEMBL web client: the **Activity client**, the **Assay client**, and the **Target client**. The connections between these clients allow access to key information for interactions.
  - **Activity client**: The extraction process began with the Activity client, which provides data on molecular activities and their associated assays and targets. The fields available in the Activity client include:

* **Target ChEMBL ID**: Identifier for the target (e.g., enzyme) in ChEMBL.
    * **Assay ChEMBL ID**: Identifier for the assay used to measure activity.
    * **Molecule ChEMBL ID**: Identifier for the molecule (e.g., inhibitor).
    * **Assay type**: ChEMBL categorizes assays into five types: Binding (B), Functional (F), ADME (A), Toxicity (T), Physicochemical (P), and Unclassified (U).
    * **Activity comment**: Active or Inactive interaction.
    * **Standard type**: For example $IC_{50}$, $K_i$, MIC, $EC_X$, $K_d$, $K_a$, and inhibition.
    * **Standard value**: Numerical value for each affinity assay.
    * **Standard unit**: In nM, uM or %
    * **SMILES**: Molecular structure in SMILES format for the small molecules.

- **Target client:** Using the retrieved **Target ChEMBL ID** from the Activity client, a connection was made to the Target client for additional target-specific data. This client provides information, including:
    * **Uniprot ID**
    * **GO ID**
    * **EC number**

- **Assay client:** The **Assay ChEMBL ID** and **Target ChEMBL ID** from the Activity client are then used to link to the Assay client, which provides assay-specific information such as:
    * **Relationship type**: this flag indicates the relationship between the target reported in the source document and the target assigned in ChEMBL:
        · **D**: Direct protein target assigned.
        · **H**: Homologous protein target assigned.
        · **M**: Molecular target other than a protein assigned.
        · **N**: Non-molecular target assigned.
        · **S**: Subcellular target assigned.
        · **U**: Default value, indicating the target has not yet been curated.
    * **Confidence score**: The confidence scores range from 0 to 9:
        · **Score 0**: Default value indicating that target assignment has yet to be curated.
        · **Score 1**: Assigned for non-molecular targets, such as cell lines or whole organisms.
        · **Score 3**: Assigned for molecular targets that are non-protein.
        · **Score 4**: Assigned when multiple homologous protein targets (e.g., a *Protein Family*) are assigned.
        · **Score 5**: Assigned when multiple direct protein targets (e.g., a *Protein Family*) are assigned.
        · **Score 6**: Assigned for homologous protein complex subunits.
        · **Score 7**: Assigned for direct protein complex subunits.

        · **Score 8**: Assigned for a homologous single protein target, often used when a direct target is not available, but a homologue from a different species is mapped. If the source of the target protein is unknown, the assignment may default to *Homo sapiens* with this score to indicate the possibility of homologue mapping.
        · **Score 9**: Assigned for a direct single protein target, reflecting the highest confidence level.

    * **Source ID**: Identifies the source or literature reference for the assay.

- **Data Mapping:**
  - UniProt ID → sequence via UniProt API.
  - Molecule ChEMBL ID → other IDs ( either PubChem CID or ChEBI ID or KEGG ID).
- **Data Filteration:**
  - Assay type = B.
  - Relationship type = D.
  - Confidence score = 9.
  - Retain only interactions with an EC number and a standard value.
  - Exclude cofactor and energy-transfer small molecules.
- **Final curated dataset:** It contains 1,407,761 enzyme-small molecule pairs.

##### S1.1.4 IUPHAR-PBS Database

- **Extraction Type:** TM.
- **Data Mining:** IUPHAR-PBS provides enzyme interaction dataset files in a comma-separated value (CSV) format, where each row contains information for one binding measurement. For each binding measurement, we extracted the PubChem SID, SMILES string for the small molecule, UniProt ID for the target, and the measured affinity. The extracted information includes:
  - **UniProt ID**
  - **PubChem SID**: PubChem SID for ligand.
  - **SMILES string**
  - **Measured affinity**: $K_i$, $IC_{50}$, $K_d$, and $EC_{50}$.
- **Data Mapping:**
  - UniProt ID → sequence via UniProt API.
  - PubChem SID → PubChem CID via PubChem API.
  - PubChem CID → SMILES string via PubChem API.
- **Data Filteration:**
  - Retain only interactions with a standard value.
  - Exclude cofactor and energy-transfer small molecules.
- **Final curated dataset:** It contains 5,076 enzyme-small molecule pairs.

##### S1.1.5 PubChem Database

- **Extraction Type:** API.
- **Data Mining:** PubChem's API does not provide an advanced filtering system comparable to ChEMBL for selecting only assay measurements related to enzymes. Given the size of the dataset (over 300 million bioactivities), downloading all assays and subsequently filtering out non-enzyme proteins would require substantial time and computational resources while also risking HTTP request limits. To address this issue, we designed a workflow that enables the extraction of maximal assay binding affinity measurements specifically involving enzymes. To achieve this, we compiled a set of tailored PubChem query terms (e.g., terms containing keywords like $IC_{50}$ or $K_d$ and others, in various combinations). We retrieved assay identifiers in batches for each term. Each assay was inspected for UniProt annotations, which were mapped to EC numbers to confirm enzymatic activity. For each query term, the process was continued until the retrieval rate of enzyme-associated assays dropped to nearly zero. At that point, we manually stopped execution and proceeded to the next term. The extracted assay IDs from each new term were cross-checked against those from previous terms to avoid redundant evaluation of enzymatic specificity. This iterative strategy ensured broad coverage of enzyme-related assays while avoiding unnecessary downloads and excessive API calls. For automatic reproducibility, it is recommended to use the provided list of assays related to the enzyme activity (`aids_with_ec_number.txt`) rather than passing all provided terms.
  - **Assay ID**: A unique identifier for each assay.
  - **Assay type**: $K_i$, $IC_{50}$, $K_d$, and $EC_{50}$.
  - **assay value**
  - **UniProt ID**
  - **PubChem SID**
  - **SMILES string**
  - **Activity comment**: Active or Inactive interaction.
  - **EC Number**
  - **Assay Description**
- **Data Mapping:**
  - UniProt ID → sequence via UniProt API
  - PubChem SID → PubChem CID via PubChem API.
  - PubChem CID → SMILES string via PubChem API.
- **Data Filteration:**
  - Retain only interactions with an EC number and a assay value.
  - Exclude cofactor and energy-transfer small molecules.
- **Final curated dataset:** It contains 481,017 enzyme-small molecule pairs.

##### S1.1.6 SABIO-RK Database

- **Extraction Type:** API.

- **Data Mining:** We accessed SABIO-RK through its RESTful web services. First, all available `EntryID`s were retrieved using a broad query (`EntryID:*`). These identifiers were then used in a POST request to export assay data in tab-delimited format, including fields such as:
  - **EntryID**: SABIO-RK unique record identifier.
  - **Standard type**: Km, Ki, $IC_{50}$.
  - **Standard value**
  - **Standard unit**
  - **UniProt ID**
  - **EC Number**
  - **Substrate name**
  - **Inhibitor name**
- **Data Mapping:**
  - UniProt ID → protein sequence.
  - Small Molecule name → PubChem ID, ChEBI ID, or KEGG ID (via PubChem API and MolToId dictionary).
  - Small molecule ID → SMILES string (via PubChem, ChEBI, or KEGG APIs).
- **Data Filteration:**
  - Retain only interactions with an EC number and a standard value.
  - Exclude cofactor and energy-transfer small molecules.
- **Final curated dataset:** It contains 4,695 enzyme-small molecule pairs.

#### S1.2 EEC-Confirmed Pairs

This category includes databases that provide enzyme–small molecule pairs associated with functional GO IDs, which can be mapped to evidence codes [1]. These evidence codes include experimental, phylogenetically inferred, computational analysis, author statement, curator statement, and electronic annotation codes. From this set, we retained only enzyme–small molecule pairs linked to experimental evidence codes (EEC), which we refer to as EEC-confirmed pairs. Such data were extracted from Brenda, Gene Ontology, Rhea, and UniProt.

##### S1.2.1 Brenda Database

- **Extraction Type:** TM.
- **Data Mining:** In the first stage, we extracted follwoing information for each catalistic activity.
  - **UniProt ID**
  - **EC number**
  - **Substrate name**
  - **Inhibitor name**
- **Data Mapping:**

- UniProt ID → sequence via UniProt API
- EC number → GO ID.
- GO-UniProt ID pair → EEC via GoToEec.
- Substrate/Inhibitor name → PubChem ID, ChEBI ID, or KEGG ID (via PubChem API and MolToId dictionary).
- Substrate/Inhibitor ID → SMILES string (via PubChem, ChEBI, or KEGG APIs).

- **Data Filteration:**
  - Retain only interactions with experimental evidence and EC number.
  - Exclude cofactor and energy-transfer small molecules.

- **Final curated dataset:** It contains 19,445 enzyme-small molecule pairs.

##### S1.2.2 Gene Ontology Database

- **Extraction Type:** TM.
- **Data Mining:** The data mining step involves extracting relevant attributes from each term in the Gene Ontology database. Each term has a format as shown below, and the following information was extracted from each term:

```
[Term]
id: GO:0003921
name: GMP synthase activity
namespace: molecular_function
def: "Catalysis of the reaction:
ATP + XMP + NH4(+) = AMP + diphosphate + GMP + 2H+." [RHEA:18301]
xref: MetaCyc:GMP-SYN-NH3-RXN
xref: RHEA:18301
is_a: GO:0016879 ! ligase activity, forming carbon-nitrogen bonds
relationship: part_of GO:
0003922 ! GMP synthase (glutamine-hydrolyzing) activity
```

  - **GO ID**
  - **Reaction ID**
  - **EC number**
  - **Substrate name**

- **Data Mapping:**
  - Reaction ID → UniProt ID (This step has been done by using cross-references file provided by Rhea website [2].)
  - Substrate name → PubChem ID, ChEBI ID, or KEGG ID (via PubChem API and MolToId dictionary).
  - Substrate ID → SMILES string (via PubChem, ChEBI, or KEGG APIs).
  - GO-UniProt ID pair → EEC via GoToEec.

- **Data Filteration:**
  - Retain only interactions with experimental evidence and EC number.

– Exclude cofactor and energy-transfer small molecules.

- **Final curated dataset:** It contains 11,729 enzyme-small molecule pairs.

##### S1.2.3 Rhea Database

- **Extraction Type:** TM.
- **Data Mining:** The data mining step involves extracting relevant attributes from each reaction in the Rhea database. Each reaction has a format as shown below and the following information was extracted from each reaction:

```
///
ENTRY        RHEA:10003
DEFINITION   H2O + pentanamide <=> NH4(+) + pentanoate
EQUATION     CHEBI:15377 + CHEBI:16459 <=> CHEBI:28938 + CHEBI
    :31011
///
ENTRY        RHEA:10004
DEFINITION   benzyl isothiocyanate = benzyl thiocyanate
EQUATION     CHEBI:17484 = CHEBI:16017
ENZYME       5.99.1.1
///
```

  – **Reaction ID**
  – **Substrate ID**
  – **EC number**

- **Data Mapping:**

  – Reaction ID → UniProt ID, this step has been done by using cross-references file provided by Rhea website [2].
  – Reaction ID → GO ID, this step has been done by using cross-references file provided by Rhea website [2].
  – GO-Uniprot ID to EEC via GoToEec.

- **Data Filteration:**

  – Retain only interactions with experimental evidence.
  – Exclude cofactor and energy-transfer small molecules.

- **Final curated dataset:** It contains 14,434 enzyme-small molecule pairs.

##### S1.2.4 UniProt Database

- **Extraction Type:** API.
- **Data Mining:** In the first step, we extracted relevant information for each Catalysis of the reaction from the UniProt database, applying specific filters to ensure data quality and relevance:

  – **Reviewed=True**: We filtered for only reviewed entries in UniProtKB/Swiss-Prot, which have been manually curated and verified by experts to ensure high-quality, reliable data.

- **Existence=1**: We included only entries with the highest level of experimental evidence, indicating that each protein has been directly observed (Protein Existence Level 1).
- **Active=True**: Only active, non-obsolete enzymes.
- **Annotation Score= 5.0**: Only enzymes with highest annotation scores.

The extracted information includes:

- **UniProt ID**
- **Sequence**
- **Reaction ID**
- **EC number**
- **Evidence code**
- **Substrate name**

- **Data Mapping:**
  - Substrate names → substrate ID (via PubChem API and MolToId dictionary).
  - Substrate IDs → SMILES (via PubChem, ChEBI, or KEGG APIs).
- **Data Filteration:**
  - Retain only interactions with experimental evidence and EC number.
  - Exclude cofactor and energy-transfer small molecules.
- **Final curated dataset:** It contains 22,927 enzyme-small molecule pairs.

### S2 Excluded Molecules

In table 1 , we list all molecules and their category for being removed. Ions and Cations: Simple ions and metal cations were excluded because their interactions are dominated by general electrostatic effects, which are highly predictable and provide little information for learning the specific molecular recognition patterns targeted by our model. Energy Transfer: Energy-transfer molecules were excluded since their interactions are highly predictable and tend to inflate model performance without reflecting true generalization. Other: Common salts, inorganic compounds, and macromolecules were excluded as they do not represent specific enzyme–small molecule binding events relevant to model learning.

### S3 EMMI dataset composition

Table 2 summarizes the contribution of each source database to the EMMI dataset after cluster-based down-sampling. The numbers do not reflect the true contribution of each database to all available enzyme–molecule interactions, since the selected interaction are effected by cluster-based down-sampling.

**Supplementary Table 1** List of small molecules excluded from the dataset.

| **Category** | **Molecules** |
|---|---|
| **Ion and Cation** | $Mn^{2+}$, $Zn^{2+}$, $Cd^{2+}$, $Cr^{3+}$, $Fe^{3+}$, $Fe^{2+}$, $Ca^{2+}$, $Mg^{2+}$, $Ba^{2+}$, $Sr^{2+}$, $Cu^{+}$, $Cu^{2+}$, $Na^{+}$, $Co^{2+}$, $Pb^{2+}$, $Rb^{+}$, $Ni^{2+}$, $K^{+}$, $Fe_2S_2{}^{2+}$, $Fe_2S_2{}^{+}$, $W^{4+}$, $Hg^{2+}$, $Se^{2-}$, $S^{2-}$, $X^{-}$, $I^{-}$, $Cl^{-}$, $Br^{-}$, $Cu^{-}$, $Ag^{+}$, $Sn^{2+}$, $Li^{+}$, $F^{-}$, $IO^{-}$, $SO_4{}^{2-}$, $CoA^{4-}$, $H^{+}$, $BrO^{-}$. |
| **Energy Transfer** | AMP, ADP, ATP, UMP, UDP, UTP, GDP, GTP, $AMP^{2-}$, $ADP^{3-}$, $ATP^{4-}$, $UMP^{-}$, $UDP^{2-}$, $UTP^{3-}$, $GDP^{2-}$, $GTP^{3-}$, AMP 3'-end$^{-}$ residue, NADH, NADP, NADPH, $NAD^{+}$, $NADP^{+}$, diphosphate$^{2-}$. |
| **Other** | NaCl, KCl,$I_2$, $NiCl_2$, $HgCl_2$, $CaCl_2$, $NiCl_2$, $CO_2$, $H_2O$, $MgCl_2$, $H_2$, CsCl, Li, LiCl, $NH_4Cl$, S, $SnCl_2$ (anhydrous), $AlF_3$, $O_2$, KBr, $CuCl_2$, Co, $AlCl_3$, $SnCl_2$, $NaBH_4$, $MnCl_2$, phosphoprotein, collagen, calmodulin, RNA, lipid, CoA, fatty acid anion, lipopolysaccharide, DNA, protein polypeptide chain. |

**Supplementary Table 2** The contribution of different databases in the EMMI dataset after cluster-based down-sampling

| Database | Enzyme—Inhibitors | Enzyme—Substrate | Enzyme—Non-interacting |
|---|---|---|---|
| Brenda | 3,020 | 4,504 | 1,869 |
| ChEMBL | 17,378 | 0 | 51,754 |
| Uniprot | 4,803 | 16,121 | 0 |
| BindingDB | 7,847 | 0 | 24,338 |
| Sabio-RK | 38 | 172 | 1,565 |
| Pubchem | 84 | 0 | 117 |
| Iuphar-BPS | 165 | 0 | 25 |
| Gobo | 0 | 6,887 | 0 |
| RHEA | 0 | 6,537 | 0 |

### S4 Degree distribution

Figure 1 illustrates the degree distributions for each split method. The red line indicates the median, and the whiskers represent the range of non-outlier values in the box plot.

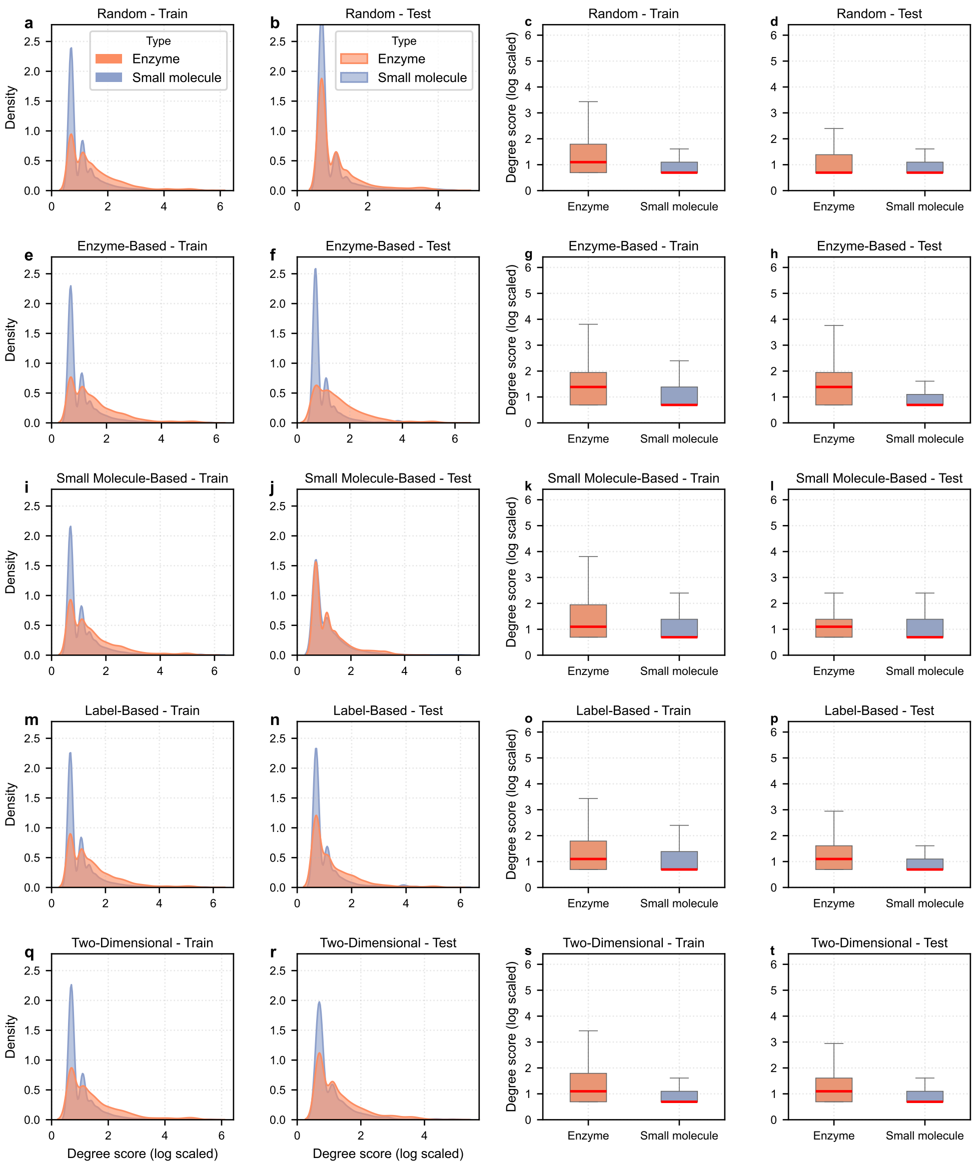


**Supplementary Figure 1** Degree distributions for each split method.
